# Complete genomic assembly of Mauritian cynomolgus macaque killer immunoglobulin-like receptor and natural killer group 2 haplotypes

**DOI:** 10.1101/2023.12.10.570943

**Authors:** Trent M. Prall, Julie A. Karl, Joshua M. Varghese, David A. Baker, Nicholas R. Minor, Muthuswamy Raveendran, R. Alan Harris, Jeffery Rogers, Roger W. Wiseman, David H. O’Connor

## Abstract

Mauritian-origin cynomolgus macaques (MCM) serve as a powerful nonhuman primate model in biomedical research due to their unique genetic homogeneity, which simplifies experimental designs. Despite their extensive use, a comprehensive understanding of crucial immune-regulating gene families, particularly killer immunoglobulin-like receptors (KIR) and natural killer group 2 (NKG2), has been hindered by the lack of detailed genomic reference assemblies. In this study, we employ advanced long-read sequencing techniques to completely assemble eight KIR and seven NKG2 genomic haplotypes, providing an extensive insight into the structural and allelic diversity of these immunoregulatory gene clusters. Leveraging these genomic resources, we prototype a strategy for genotyping KIR and NKG2 using short-read, whole exome capture data, illustrating the potential for cost-effective multi-locus genotyping at colony scale. These results mark a significant enhancement for biomedical research in MCMs and underscores the feasibility of broad-scale genetic investigations.

## Introduction

Rhesus (*Macaca mulatta*) and cynomolgus (*Macaca fascicularis*) macaques share a close evolutionary history with humans. Due to their similar immune systems, macaques are commonly employed as biomedical models of human health (1–3). Whereas rhesus macaques have been traditionally favored, the popularity of cynomolgus macaques has increased due to the limited supply of rhesus following an export ban in 1978. Furthermore, due to the recent prohibition of macaque exports from China and the current investigations into the illegal trade of Cambodian monkeys, Mauritius has emerged as the primary exporter of nonhuman primates (NHP) (4, 5). Mauritian cynomolgus macaques (MCM) are unique as they descend from a small founding population and harbor restricted genetic diversity (6). This limited diversity makes MCM particularly valuable in basic biologic and pharmacogenetic experiments that benefit from tight genetic control of experimental cohorts. Genetically-defined MCM have been used to model simian immunodeficiency virus, tuberculosis, SARS-CoV-2, transplantation, and other disease and pharmacokinetic contexts (7–12).

Despite their widespread use in research, the cynomolgus macaque draft genome contains notable gaps and omissions (13, 14). This incompleteness hinders robust analyses, given the pivotal role of high-quality reference genomes in biological research (15). High-quality genomic references ensure sequencing reads from experimental samples can be reliably mapped, facilitating the generation of comprehensive annotation of genomic features. Notably, genomic characterizations provide a broader perspective than transcript sequencing alone, as the latter, influenced by tissue-specific transcriptional profiles, might not fully represent the underlying genetic architecture (16–18). Robust annotations are pivotal for precision in functional genomics and transcriptomics. Furthermore, high-quality genomic data facilitate the creation of large-scale genotyping assays, essential for the characterization of experimental animals for a more comprehensive understanding of experimental outcomes. Thus, refining and completing the cynomolgus macaque genome is imperative for advancing our understanding and harnessing this model organism’s full potential in biomedical research.

Gaps in the cynomolgus draft genome are commonly present within immune receptor gene families. This is primarily due to the high saturation of short tandem repeats, homopolymer stretches, and multi-copy gene clusters within these regions (19). For example, the major histocompatibility (*MHC*) genes, known as the human leukocyte antigens (*HLA*) in humans, are considered the most polymorphic gene cluster of mammalian genomes (20). *HLA* haplotypes contain three highly polymorphic class I genes: *HLA-A, HLA-B, and HLA-C*. In contrast, the *MHC* region of macaques has undergone intricate duplications, deletions, and rearrangements such that each haplotype has gained variable numbers of less polymorphic *MHC-A* and *MHC-B* loci (21–23). A similar level of genetic complexity extends into *MHC*-related gene families. For instance, the killer immunoglobulin-like receptors (*KIR*) recognize epitopes within the three-dimensional structures of classical MHC class I proteins (24). Similarly, the natural killer group 2 (NKG2) receptors (encoded by the killer cell lectin receptors, *KLR*, genes) heterodimerize with CD94 and target a non-classical MHC class I molecule, MHC-E (25). The co-evolutionary dynamics shared among these interacting families have driven their expansion in macaques relative to humans (26). The human *KIR* genes are categorized into four lineages based on MHC binding specificity and phylogenetic relationships (27). Lineage III *KIR*, specific for *HLA-C*, displays increased diversity in humans. Since the *HLA-C* locus is believed to have been duplicated from *HLA-B* after the divergence of humans and macaques from their most recent common ancestor (28), the absence of an *HLA-C* ortholog in macaques has resulted in an expanded lineage II *KIR* gene repertoire that is specific to MHC-A and MHC-B ligands (29). Similarly, *NKG2C* has duplicated from one to three gene copies in macaques, mirroring the duplication of *MHC-E* in macaques relative to humans (30). These clusters of highly related genomic sequences require careful attention to resolve and catalog properly.

Resolving the *MHC*, *KIR*, and *NKG2* genomic regions is difficult due to the limitations of short-read sequencing technologies. These limitations can be circumvented by single-molecule sequencing platforms such as Pacific Biosciences (PacBio) and Oxford Nanopore Technologies (ONT) that can generate reads spanning hundreds of kilobase pairs (31). Reads of substantial length unambiguously span repetitive genomic elements allowing continuous, chromosomal-scale de novo assemblies to be accurately resolved (32). Recently, long-read techniques have been used to resolve *MHC* and *KIR* genomic haplotypes from macaques. We previously described a comprehensive five-megabase pair (Mb) MCM *MHC* genomic assembly using a hybrid sequencing technique (22). In that effort, we combined sequencing technologies, capitalizing on the structural insights of ONT ultra-long reads, and the precision of PacBio high fidelity (HiFi) reads to define the MHC region accurately. Another group resolved two *MHC* and two *KIR* haplotypes from a Vietnamese-origin cynomolgus macaque using PacBio HiFi (23). Yet another group resolved six *KIR* haplotypes from rhesus macaques using CRISPR/Cas9 enrichment followed by ONT sequencing ((30, 33)). The *NKG2* region has remained less well characterized with only a single *NKG2* haplotype from rhesus macaque described from bacterial artificial chromosomes (30, 33). Nevertheless, great care was taken in all of these examples to manually validate annotations to guarantee accurate representations of the genomic content within each haplotype. In contrast, the *KIR* and *NKG2* genomic regions in MCM have not been fully characterized to date.

Amplicon sequencing experiments with MCM have shown that seven and eight haplotypes encompass *MHC* and *KIR* genetic diversity, respectively (34–37). Because the *MHC* and *KIR* are encoded on separate chromosomes, it can be reasoned that the pangenome of MCM displays a similar level of diversity. It is therefore, theoretically possible to establish genetic control over multiple loci when selecting experimental cohorts. However, this requires the development of comprehensive characterizations of the key immune-related gene families to develop high throughput and cost-effective genotyping strategies. As mentioned earlier, we previously developed a hybrid, long-read assembly approach to resolve an MCM *MHC* region (22). Here, we extend this methodology to assemble *KIR* and *NKG2* genomic regions from thirteen representative MCM individuals. Employing this approach, we provide a thorough genomic characterization of the eight known *KIR* haplotypes and introduce seven new *NKG2* haplotypes. Additionally, we leverage the accuracy of the resulting genomic assemblies to prototype an Illumina-based, exome target-capture approach for multi-locus genotyping from a single experiment. Overall, our findings give a detailed insight into the genetic variance within these critical immune receptor regions, underscoring their relevance in MCM-oriented biomedical research.

## Materials and Methods

### Animal selection

Peripheral blood mononuclear cells (PBMC) and splenocytes were obtained from thirteen MCM housed at the Wisconsin National Primate Research Center during semi annual health checks or at necropsy. Animals were selected for analysis based on previously established *MHC* and *KIR* genotypes (34, 37–39). Sampling was performed in concordance with protocols approved by the University of Wisconsin-Madison Institutional Animal Care and Use Committee as well as guidelines contained within the Animal Welfare Act, the Guide for Care and Use of Laboratory Animals, and the Weatherall report (40).

### High molecular weight DNA isolation

High molecular weight (HMW) DNA was prepared as previously described (22). DNA was extracted from ~5×10^6^ PBMCs using New England Biolab’s Monarch high-molecular-weight DNA extraction kit, following the manufacturer’s protocol. PBMCs were pelleted at 1000 g for 3 minutes, resuspended in 150 μL nuclei prep solution, and mixed by pipetting 10 times. Next, 150 μL of nuclei lysis solution was added, inverted 10 times, and incubated for 10 minutes at 56°C at 300 rpm for ONT libraries and 2000 rpm for PacBio libraries. After adding 75 μL precipitation buffer and inverting 10 times, two DNA capture beads and 275 μL isopropanol were added, followed by 8 minutes of vertical rotation at 10 rpm.

Supernatants were removed from samples, avoiding disrupting gDNA on capture beads. Two washes were performed using 500 μL DNA wash buffer, inverting three times, and carefully removing the wash buffer. Beads were transferred to a bead retainer, and pulse spun for ~1 second. Then, 100 μL elution buffer was added to a 2-mL microfuge tube containing separated glass beads and incubated for 5 minutes at 56°C at 300 rpm. The eluate was separated from DNA capture beads using the supplied bead retainer and transferred to an Eppendorf DNA LoBind 1.5-mL tube. The bead retainer and tube were centrifuged at 12,000g for 1 minute, and the final eluate with DNA was stored at 4°C until library preparation.

### Preparation of ultra-long DNA libraries for ONT sequencing

Sequencing libraries were prepared with ONT SQK-RAD004 rapid sequencing kits using a previously described method (31). This method employs a robotic pipette to combine sequencing reagents with HMW DNA by pipetting as slowly as possible. These measures decrease DNA shearing and help to maintain ultra-long read lengths. For each library, 1.5 μl FRA and 3.5 μl elution buffer were added to a 16 μl DNA aliquot, mixed by pipetting five times, and incubated at 30°C for 1 minute, followed by 80°C for 1 minute on an Applied Biosystems Thermal Cycler. Next, 1 μl RAP was added to the solution and pipetted five times. The library was then incubated at room temperature for approximately 10 minutes while flow cells were primed. After priming, 34 μl SQB and 20 μl water were added to the sample solution and pipetted three times.

For priming, 30 μl FLT was added to a tube of FLB, and a small volume of buffer was removed from the priming port. Then, 800 μl of FLT + FLB solution was added to the priming port, and after a 5-min incubation, 200 μl of FLT + FLB solution was slowly added to the priming port, allowing a small volume to rise from the SpotOn port and return to the cell. 75 μl of the prepared library was slowly drawn into a pipette with a wide bore tip. The library was added dropwise to the SpotON port.

### ONT sequencing and base calling

Twelve to sixty ultra-long libraries were sequenced from each selected animal (Supplemental Table I). Libraries were sequenced using R9.4 (FLO-MIN106) flow cells according to ONT guidelines. Flow cells were sequenced on a GridION instrument with the installed MinKnow software. Multiple versions of MinKNOW were used through the duration of the study as updates were released and specific versions were recorded in the metadata of FAST5 files generated for each run. After 18 hours, sequencing was paused, and the flow cells were flushed using EXP-WSH004 flow cell wash kits. Flow cells were then re-primed and loaded following the same procedure described before. After an additional 24 hours, sequencing was paused again, and the flow cells were washed, primed, and reloaded. After a second library reload, each flow cell was run until all pores were exhausted. The raw FAST5 data for each run were merged into a single FAST5 per animal. Base calling was performed on merged FAST5 files using Bonito version 0.3.8 on A100 GPU hardware running CUDA 11.2.

### PacBio HiFi sequencing

High-molecular-weight DNA for each sample was provided to the University of Wisconsin–Madison Biotechnology Center DNA Sequencing Facility. The DNA quality was measured using a Thermo Fisher Scientific NanoDrop One instrument, recording concentrations, 260/230 ratios, and 260/280 ratios. The extracted DNA was quantified with the Thermo Fisher Scientific Qubit dsDNA high-sensitivity kit, and samples were diluted before analysis on an Agilent FemtoPulse system to assess DNA sizing and quality.

PacBio HiFi libraries were prepared following PN 101-853-100 version 03 (PacBio) protocol, including modifications such as shearing with Covaris gTUBEs and size selection using Sage Sciences BluePippin. Library quality was assessed with the Agilent FemtoPulse system, and the library was quantified using the Qubit dsDNA high-sensitivity kit. Sequencing was performed on a PacBio Sequel II instrument with the Sequel Polymerase Binding Kit 2.2 at the University of Wisconsin–Madison Biotechnology Center DNA Sequencing Facility. Raw sequencing data were converted to circular consensus sequencing (CCS) FASTQ files using SMRT Link version 8.0.

### De novo assembly of KIR and NKG2 regions

ONT and PacBio HiFi FASTQ reads from each sample were mapped to the human reference genome GRCh38 (GCA_000001405) using minimap2 version 2.17 (41) with flags “-ax map-ont” and “-ax map-hifi.”. Reads that aligned to genomic coordinates Chr19:54,014,000-55,240,000 (VSTM1 - TMEM868) and Chr12:9,090,000-11,425,000 (M6PR - PRB3) were extracted using SAMtools version 1.11 (42) for de novo assembly of the *KIR* and *NKG2* regions, respectively. After extraction, reads were converted from BAM to FASTQ format using BBTools reformat.sh https://sourceforge.net/ projects/bbmap/). Extracted ONT and PacBio FASTQ reads were assembled using hifiasm v0.19.2-r560 with the flag “--ul” used for ONT integration (43, 44).

### Error correction of genomic assemblies

Error correction was carried out on extracted PacBio HiFi reads aligning to human coordinates for *KIR* or *NKG2* using Geneious Prime v 2023.0.4 (https://www.geneious.com/). The Geneious mapper was employed with custom sensitivity and up to 10 iterations for fine-tuning. Several advanced custom settings were enabled: map multiple best matches set to none; trim paired read overhangs; map only paired reads that both map; allow a maximum of 10% gaps per read with a maximum gap size of 15, word length of 18, and index word length of 13; ignore words repeated more than 12 times; allow a maximum of 5% mismatches per read; set maximum ambiguity to four; and accurately map reads with errors to repeat regions.

A consensus sequence was generated in Geneious Prime using a 0% (majority) threshold. The reference (hybrid scaffold) was called if coverage was less than five reads or if there was no coverage. The consensus sequence was corrected to match higher-accuracy PacBio HiFi reads at positions where PacBio data differed from the hybrid assembly, provided PacBio coverage was greater than five reads. Some regions in the hybrid assembly lacked PacBio coverage and remained uncorrected, likely corresponding to areas where the macaque differed significantly from the corresponding human genomic region. To address these areas, a second round of PacBio mapping using minimap2 was performed, and reads aligning to the first-round error-corrected hybrid assembly were extracted. This new set of PacBio reads was used for a second round of error correction and to fill gaps left by the initial PacBio reads. The final mapping yielded end-to-end, >5X PacBio coverage across the entirety of all assemblies.

### Gene annotation

We utilized Exonerate version 2.4.0 with the “est2genome” mapping model for sequence comparison (https://www.ebi.ac.uk/about/vertebrate-genomics/software/exonerate). Human gene and CDS annotations taken from the GRCh38 *KIR* or *NKG2* genomic regions were used to run Exonerate recursively. Results were filtered for matches >95%, and the corresponding annotation tracks were loaded onto the error-corrected assemblies in Geneious Prime. Annotations were manually curated to retain a single gene annotation per locus, as well as CDS as appropriate. All annotated genes were individually compared against available human and rhesus macaque orthologs to confirm proper annotation. Gene names were assigned based on human and rhesus macaque orthologs.

### Generation of genomic figures

Genomic annotation maps were generated from GFF3 files exported from Geneious Prime with the karyoploteR package (45) within RStudio (RStudio Team 2022; https://www.posit.co). Any additional formatting to improve legibility was performed with Adobe Illustrator.

### NKG2 allele nomenclature

*NKG2* allele nomenclature was assigned to based on established conventions used for NHP KIR nomenclature (46, 47). Briefly, genes were characterized using a two-digit numerical sequence, with non-synonymous allele variations indicated by a three-digit number following an asterisk. Synonymous changes within the coding sequence are further specified by an additional two-digit number after a colon. Additionally, intron substitutions are identified by a third set of digits, also separated by a colon, placed after the synonymous variant number.

### Whole exome sequencing

PBMC and/or splenocytes from 85 MCM were chosen for whole exome sequencing. Genomic DNA was isolated on a Maxwell RSC 48 robot with Maxwell RSC buffy coat DNA kits (Promega). Isolated DNA was analyzed for purity and molecular weight using PicoGreen and gel imaging. After DNA quality control tests, Illumina sequencing libraries with incorporated barcodes were produced following standard procedures (48) using human genome sequencing center custom exome design (HG38_HGSC_Twist_Comprehensive_Exome) and Rhesus Spike-In probes following manufacturer’s protocol (https://www.twistbioscience.com/). This new design is a modification of our previous Rhexome design using NimbleGen Human whole exome probes plus rhesus-specific probes (49). Groups of ten macaque bar-coded samples were pooled and captured together (10Plex). Seven resulting pools of ten samples each, enriched for the macaque exome by the capture process, were sequenced in a single lane of an Illumina NovaSeq instrument. This procedure results in an estimated sequence read depth greater than 20X for 99% of on-target reads.

### Genotyping from exome data

We developed a custom pipeline to evaluate the potential of extracting *KIR* and *NKG2* genotype information from exome data generated using human capture probe sets, available at (https://github.com/dholab/iWES-genotyper). This tool dynamically selects regions within each allele expected to provide adequate coverage based on the observed coverage from all samples for a specific allele. To begin, FASTQ files from all exome datasets are mapped to a comprehensive reference library containing every gDNA sequence identified from the haplotype assemblies. Subsequently, the depth of coverage for each allele across all samples is superimposed. Nucleotide positions within the gDNA segment achieving a cumulative depth of coverage of 30 or more are deemed necessary for the individual depth of coverage threshold to make a positive call. Conversely, positions falling short of the 30-depth threshold are “masked” and are not mandatory for positive identifications. The coverage profiles for individual samples are then cross-referenced against the masked gDNA allele library to determine positive calls. An allele is marked positive for a particular sample if its coverage depth surpasses 3 across all mandatory positions. Finally, these data plots are transformed into a summary report indicating the median depth of coverage over all necessary positions.

## Results

### Genomic assembly of eight MCM KIR haplotypes

Thirteen representative MCM were chosen for analysis based on previous genotyping results such that each of the eight known *KIR* haplotypes was represented by at least one animal (Supplemental Table I). We performed whole genome sequencing using ONT and PacBio DNA libraries on individual animals. From these data, all eight genomic haplotypes were completely assembled from the centromeric *LILRA6* to the telomeric *NCR1* loci that flank the *KIR* gene cluster (Figure 1). The assemblies ranged significantly in length, with the shortest K4 haplotype spanning 115,859 bp and the longest K5 haplotype spanning 199,303 bp measured from the start codon of the centromeric *KIR3DL20* to the stop codon of the most telomeric *KIR3D* locus. We previously characterized forty-nine *Mafa-KIR* transcripts by transcript sequencing (34, 37). To assess the nucleotide-level accuracy of each haplotype assembly, we aligned these previously characterized *KIR* transcripts against their respective haplotype. All forty-nine transcripts aligned with perfect identity across exon intervals. Therefore, we believe these assemblies depict near-perfect resolutions of the genomic nucleotide sequence.

**Figure 1.**
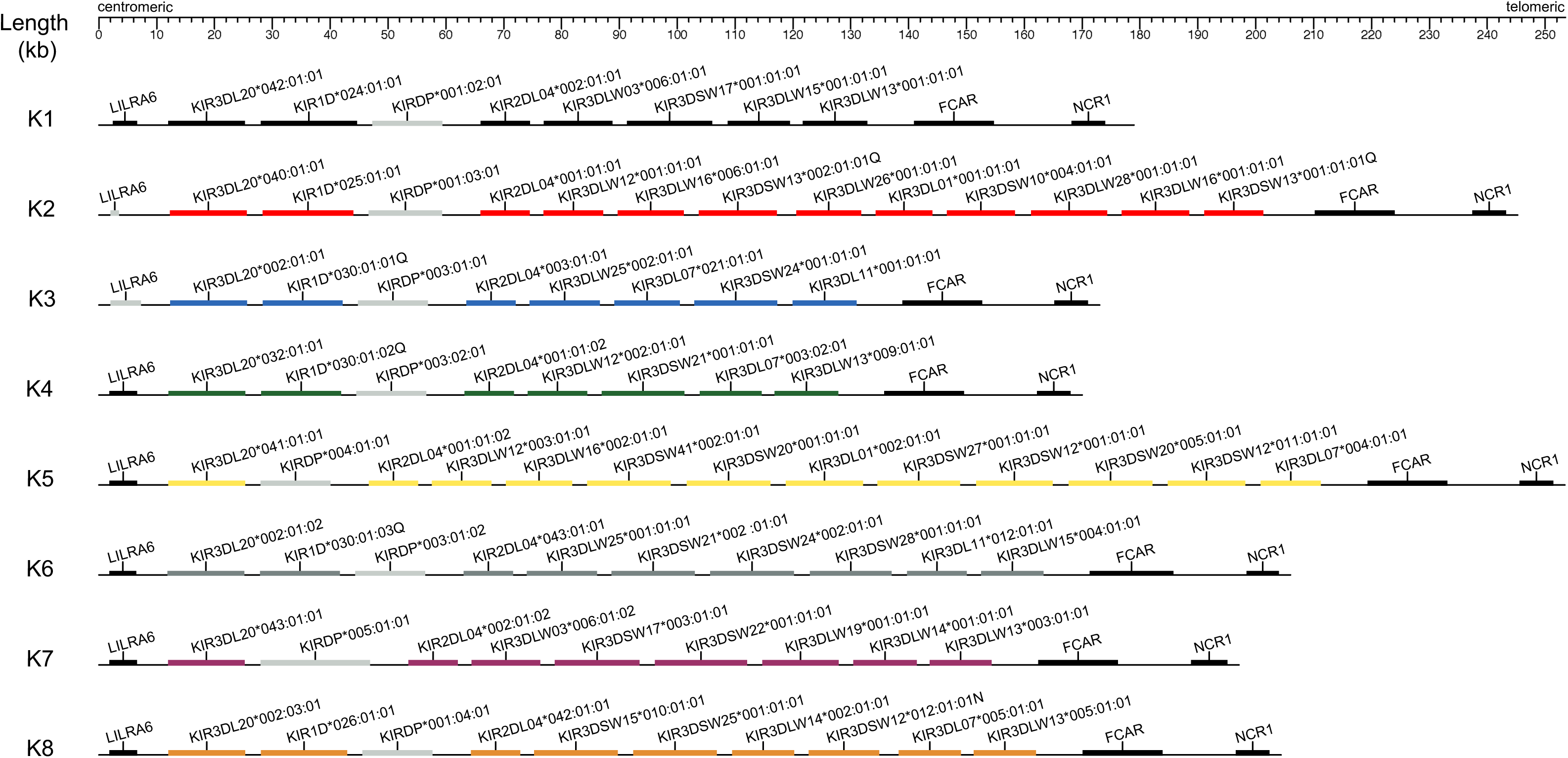
Gene content of eight *KIR* haplotypes of the MCM genomic region. The KIR genomic region is flanked by the centromeric *LILRA6* and telomeric *FCAR* and *NCR1* genes. Flanking genes are displayed in black. *KIR* genes are depicted by colored boxes that match the corresponding haplotype. Noncoding pseudogenes are depicted with grey boxes. The position in kilobases is displayed at the top of the figure.

We identified twenty-one *KIR* genes that were not identified by previous transcript sequencing. Of these newly identified genes, fourteen represented novel alleles that have not been characterized in other cynomolgus macaque populations. We submitted novel sequences to the Immuno Polymorphism Database (IPD) for official allele designations (Supplemental Table II) (47). The novel genes encompassed five *KIR3DL20*, two *KIR2DL04*, five *KIR3DS*, and four *KIR3DL*. The majority of the undetected transcripts can be attributed to sequence mismatches at primer binding sites. The remaining unaccounted-for genes might have been overlooked because each lymphocyte subpopulation expresses only a subset of the total *KIR* repertoire encoded within the genome (50). These results increase the range of *KIR* genes encoded per haplotype to seven to twelve within the MCM population (Table I). This is somewhat larger than other cynomolgus macaque populations which are estimated to encode three to thirteen genes per haplotype (51). This may be the result of a founder effect in which gene-scarce haplotypes were absent from the founding population. However, prior characterizations of haplotypes in non-Mauritian animals were conducted using transcript sequencing, which, as our current data suggests, could potentially lead to an underestimation of the range of genes actually encoded.

**Table I.**
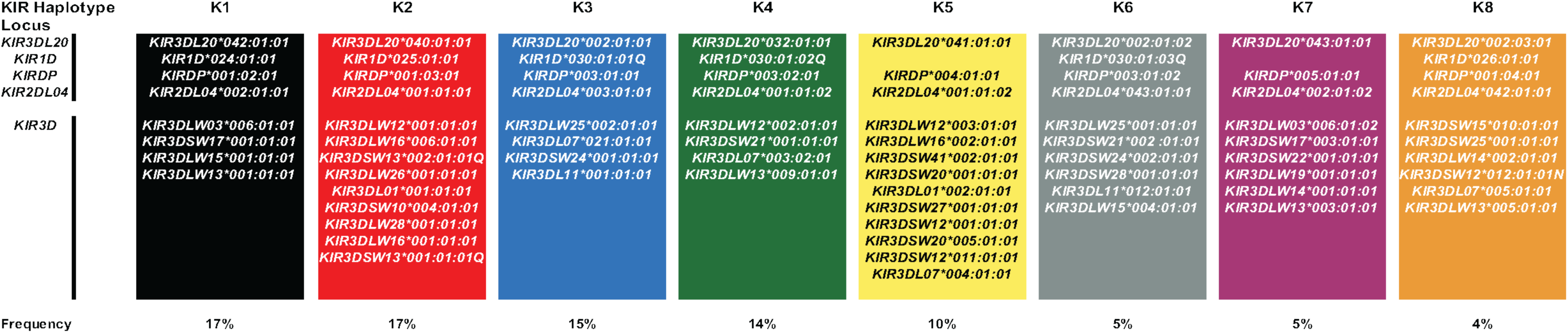
*KIR* alleles per MCM haplotype. Alleles are displayed in alphabetical order by gene and not in chromosomal order. Allele nomenclature was classified by IPD. The estimated population frequency of each haplotype is displayed below each haplotype (34).

### Centromeric KIR loci are preserved between haplotypes

The centromeric region of macaque *KIR* haplotypes includes three expressed loci: *KIR3DL20*, *KIR1D*, and *KIR2DL04*. To date, no ligands have been identified for these genes; however, it is speculated that they may bind more conserved ligands such as non-classical MHC class I proteins due to their high prevalence within the species. Our assemblies uncovered a *KIR3DL20* gene on all eight haplotypes, seven of which were previously unknown. These newly identified genes included five novel alleles that were not recovered by transcript sequencing due to primer mismatches. *KIR3DL20* is evolutionarily distinct from other *KIR* loci within macaques (52). Thus, the flanking untranslated region (UTR) of these newly identified *KIR3DL20* loci significantly differ from the limited *KIR3D* UTR genomic sequences used to design our previous primer set, resulting in inefficient binding. This highlights the importance of comprehensive genomic data when developing genetic technology. We observed the highest level of allelic variation in *KIR3DL20* with five distinct alleles. This level of diversity is also observed in non-Mauritian populations (51). This study also suggested that the K3 haplotype is shared between Mauritian and Malaysian-origin animals (designated as Cy-H9 haplotype in non-Mauritian animals) despite K3 lacking a *KIR3DL20*002* locus (37). Our new data confirms that the *KIR3DL20*002* allele is indeed shared by the K3 and Cy-H9 haplotypes. However, we also uncovered a novel *KIR3DL07*021:01:01* on the K3 haplotype that was not characterized on the Cy-H9 haplotype. Whether this gene is specific to MCM or was missed by previous transcript sequencing efforts will require additional analysis. Once again, this observation highlights the advantage of genomic assembly over transcript sequencing for haplotype characterizations. Regardless, our assemblies confirm the framework status of *KIR3DL20* in MCM, consistent with this designation in other macaque populations.

KIR2DL04 is the only KIR in macaques with a direct human ortholog. In humans, KIR2DL4 binds non-classical HLA-G, but given the lack of a functional *MHC-G* gene in macaques and the presence of hybrid MHC-AG molecules, there is speculation that KIR2DL04 may bind *MHC-AG* to perform a similar biological function. We identified two novel *KIR2DL04* genes on the K6 and K8 haplotypes. Again, these novel alleles were missed due to nucleotide mismatches within the binding site of the previously used *KIR2DL04*-specific primer set. These novel genes elevate the *KIR2DL04* to framework status within the MCM population. This is consistent with its existence on 94% of haplotypes characterized in non-Mauritian cynomolgus macaques and its framework status in humans (51)(53)(51). We observed five distinct *KIR2DL04* alleles across eight haplotypes. This mirrors the high level of *KIR2DL04* allelic diversity in other cynomolgus populations. Whether *KIR2DL04* performs an evolutionarily conserved function in macaques remains speculative despite its high prevalence within the genome.

Many macaque haplotypes contain a *KIR1D* gene of unknown function as it contains only a single immunoglobulin domain and lacks a cytoplasmic tail (54). We confirmed *KIR1D* genes on six of the eight haplotypes. We cataloged genomic extensions of previously characterized *KIR1D*024*, *KIR1D*025*, and *KIR1D*026* alleles on the K1, K2, and K8 haplotypes, respectively. In addition, we found that KIR1D*030Q is shared between the K3, K4, and K6 haplotypes. *KIR1D*030Q* has also been observed within Indonesian and Malaysian cynomolgus populations. Transcript analysis has revealed that *KIR1D*030Q* lacks an immunoglobulin domain, casting further doubt on its functional capabilities compared to other *KIR1D* variants. Our data revealed that all three *KIR1D*030Q* loci contain a large, ~1,500 bp deletion spanning part of the second intron and into exon three. This deletion removes the canonical splice acceptor site 5’ of exon three, thereby skipping its integration into the expressed mRNA. Therefore prior detections of this allele were not merely documenting a splice variant; indeed, the canonical *KIR1D*030Q* open reading frame genuinely lacks the immunoglobulin domain. We did not identify *KIR1D* transcripts from the K5 and K7 haplotypes in our previous sequencing efforts. K5 appears to have undergone a large deletion event that completely removed the *KIR1D* gene from this haplotype. K7 contains a fusion pseudogene in which the head of the ancestral *KIR1D* gene is fused with the tail of a *KIRDP* pseudogene, resulting in a non-functional CDS.

### Diversification of telomeric KIR loci

The telomeric region of macaque *KIR* haplotypes is populated by variable numbers of *KIR3DL/S* genes specific for *MHC* class I A and B ligands (52). Our eight assemblies ranged from one to six *KIR3DS* and three to five *KIR3DL* genes. In total, we identified twenty-four distinct *KIR3DS* and *KIR3DL* genes within the population. The highest level of allelic polymorphism was observed for *KIR3DLW13* and *KIR3DL07*, followed by *KIR3DLW12*, *KIR3DLW15*, *KIR3DLW16*, and *KIR3DSW12* due to their appearance on multiple haplotypes. *KIR3DL07* shares an ortholog with rhesus macaques and harbors high levels of allelic polymorphism in non-Mauritian populations (51). This suggests that *KIR3DL07* may perform some conserved or advantageous biological function, though the nature of that function is still elusive. The remaining genes with higher allelic diversity only display increased allelic diversity within the Mauritian population. This highlights the unparalleled diversification of telomeric *KIR* genes given that the MCM population was geographically isolated from southeast Asia only ~500 years ago.

We observed only one instance of allelic variants for the telomeric *KIR3DL/S* genes being shared by haplotypes, *KIR3DL28*001:01:01*, encoded on both K2 and K6. The two largest haplotypes, K2 and K5, contained multiple copies of the same *KIR3DL/S* genes. K2 encoded two copies of *KIR3DSW13* and *KIR3DLW16*, while K5 encoded two copies of *KIR3DSW12* and *KIR3DSW20*. The duplicated genes on K2 appear to be the result of a larger duplication event, as KIR3DSW16 and *KIR3DSW13* are adjacent to one another at both locations on the assembly. The K5 haplotype duplications are the result of a more complex structural event as both KIR3DSW12 genes flank the second *KIR3DSW20* towards the telomeric end of the cluster. Additional clusters of genes appearing in the same order are shared between haplotypes. For example, *KIR2DL04*002*, *KIR3DLW03*006*, and *KIR3DSW17* are shared in the same orientation between K1 and K7. Similarly *KIR2DL04*001* and *KIR3DLW12* are shared, in order, between K2, K4, and K5.

We can categorize the eight haplotypes into two general groups based on gene content. The first group (K1, K3, K4) contains a single *KIR3DS* and three *KIR3DL* genes. K1 and K4 appear to be the most closely related in that they share the positional order of genes on the chromosome, and both contain *KIR3DLW13* in their most telomeric position. This arrangement of genes is not consistent in the K3 haplotype despite K3 containing the same gene content. Group two haplotypes (K2, K5, K6, K7, K8) harbor variable gene copy numbers and increased allelic polymorphism. Within this group, three haplotypes (K6, K7, K8) contain six total KIR3D genes. The order and type of these genes are highly variable between one another. K2 and K5 contain expanded telomeric regions with nine and ten *KIR3D* genes, respectively.

### Genomic assembly of seven MCM NKG2 haplotypes

Primate immune cells employ NKG2 to measure cell surface expression of MHC class I proteins on target somatic cells (55). The *NKG2* region displays less genetic heterogeneity than *KIR*, though species-specific expansions within *NKG2* genes have been documented suggesting that it may be subject to similar evolutionary pressures as *KIR* (30). The *NKG2* region within cynomolgus macaque reference genomes are inaccurate both in contiguousness and annotation (14). To improve our understanding of these genes, we assembled the NKG2 genomic region from the thirteen long-read datasets generated during this study. This analysis revealed seven distinct haplotypes (Figure 2). This level of haplotype diversity aligns with findings in other immune regions of the MCM population (36). Therefore, the haplotypes we resolve likely encompasses the majority of genetic diversity within the population. However, one or more rare haplotypes may have been missed as *NKG2* genotypes were not screened when selecting animals evaluated in this study. Nevertheless, our work significantly expands the available *NKG2* genomic data for MCM.

**Figure 2.**
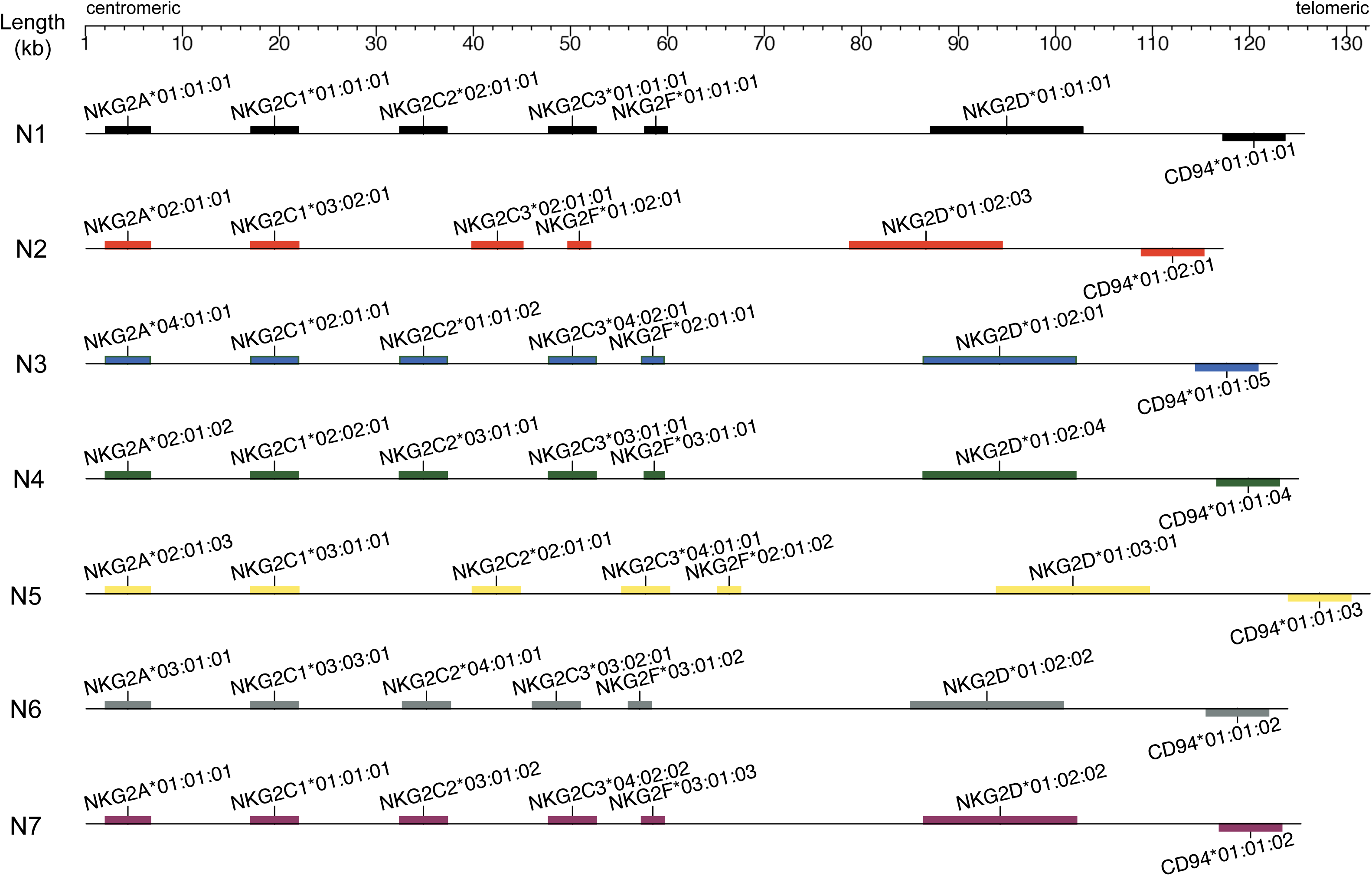
Gene composition of seven *NKG2* haplotypes in the MCM genomic region. The NKG2 genomic region stretches from the centromeric *NKG2A* and telomeric *CD94* genes. Genes on the top of the scaffold are coded in the positive sense, whereas genes on the bottom are coded in the negative sense. The position in kilobases is displayed at the top of the figure.

The genomic assemblies each contained *NKG2A*, multiple *NKG2C*, *NKG2F* and *NKG2D* genes arranged in a head to tail configuration beginning at the centromeric end of the haplotype, as well as a *CD94* open reading frame arranged in the opposite orientation at the telomeric end. The haplotypes ranged from 115,175 to 130,326 bp in length, marked from the *NKG2A* start codon to the *CD94* stop codon. The largest structural difference between haplotypes was due to insertion of an ~9,000 bp endogenous retrovirus observed within the N2 and N5 assemblies between the first and second *NKG2C* loci. Further studies are required to determine if any functional significance results from this endogenous retrovirus. Interestingly, the N2 haplotype contains a ~15,000 bp deletion, completely removing the *NKG2C2* gene. To our knowledge, this is the first observation of copy number variation within the *NKG2C* genes of macaques. The N2 haplotype was observed in three of the thirteen sequenced animals, suggesting that it may be present at a higher frequency within the MCM population. The functional consequence of this deletion is currently unknown and will require future experimentation.

### Characterization of NKG2 alleles

We employed phylogenetic analysis based on both DNA and protein sequences to systematically categorize *NKG2* alleles. The *NKG2* alleles were named adhering to the conventions of IPD’s nonhuman primate KIR database (47). In short, allelic lineages were defined by three or more unique nonsynonymous variants followed by a first colon delimiter denoting synonymous variants and a second delimiter denoting intronic variants. We characterized allelic lineages for four *NKG2A*, three *NKG2C1*, three *NKG2C2*, four *NKG2C3*, and three *NKG2F* family members. The *NKG2D* and *CD94* genes were limited to single lineages within the current collection of haplotypes (Table II). The *NKG2* genes display less allelic polymorphism per locus than the KIR genes. For example, the activating *NKG2C* genes contained only ten total allelic lineages whereas nineteen activating *KIR3DS* lineages were observed in the MCM population. The overall trend of decreased heterogeneity is consistent between primate species (30). This may be the consequence of *NKG2* molecules binding more-conserved ligands than their KIR counterparts.

**Table II.** *NKG2* alleles per MCM haplotype. *NKG2* allele nomenclature was determined following IPD’s nomenclature scheme used to name nonhuman *KIR* alleles (47). The estimated population frequency is displayed below each haplotype.

| NKG2 Haplotype Locus | N1 | N2 | N3 | N4 | N5 | N6 | N7 |
| --- | --- | --- | --- | --- | --- | --- | --- |
| NKG2A | NKG2A*01:01:01 | NKG2A*02:01:01 | NKG2A*04:01:01 | NKG2A*02:01:02 | NKG2A*02:01:03 | NKG2A*03:01:01 | NKG2A*01:01:01 |
| NKG2C | NKG2C1*01:01:01<br>NKG2C2*01:01:01<br>NKG2C3*01:01:01 | NKG2C1*03:02:01<br>NKG2C3*02:01:01 | NKG2C1*02:01:01<br>NKG2C2*01:01:02<br>NKG2C3*04:02:01 | NKG2C1*02:02:01<br>NKG2C2*03:01:01<br>NKG2C3*03:01:01 | NKG2C1*03:01:01<br>NKG2C2*02:01:01<br>NKG2C3*04:01:01 | NKG2C1*03:03:01<br>NKG2C2*04:01:01<br>NKG2C3*03:02:01 | NKG2C1*01:01:01<br>NKG2C2*03:01:02<br>NKG2C3*04:02:02 |
| NKG2F | NKG2F*01:01:01 | NKG2F*01:02:01 | NKG2F*02:01:01 | NKG2F*03:01:01 | NKG2F*02:01:02 | NKG2F*03:01:02 | NKG2F*03:01:03 |
| NKG2D | NKG2D*01:01:01 | NKG2D*01:02:03 | NKG2D*01:02:01 | NKG2D*01:02:04 | NKG2D*01:03:01 | NKG2D*01:02:02 | NKG2D*01:02:02 |
| CD94 | CD94*01:01:01 | CD94*01:02:01 | CD94*01:01:05 | CD94*01:01:04 | CD94*01:01:03 | CD94*01:01:02 | CD94*01:01:02 |
| Frequency | 42% | 20% | 9% | 6% | 14% | 2% | 5% |

### Genotyping KIR and NKG2 from whole exome data

Utilization of the restricted genetics of MCM would benefit from the ability to simultaneously screen multiple loci for desirable genotypes and/or haplotypes. Performing long-read genomic assembly or full-length gene amplification is cost-prohibitive at the colony level. To alleviate the cost, human capture probe arrays have been used to sequence exomes from rhesus and cynomolgus macaques to varying success (37, 56, 57). We previously performed *MHC* genotyping from rhesus macaque exome data, though the probe design included a collection of probes designed to specifically enrich rhesus macaque *MHC* sequences (49). This method requires sufficient read coverage over predetermined diagnostic regions to identify each allele. The efficiency of cross-species target capture is variable without probes tailored for the genome’s divergent sequences. Also, efficiently captured regions in the genomes are discontinuous, often necessitating multiple separate regions for an accurate allele determination. To address these issues, we developed a pipeline that adjusts diagnostic subregions within each allele based on the median read coverage across all pooled samples, accommodating uneven coverage across exon intervals resulting from inefficient target capture (see methods).

We sequenced eighty-five MCM exomes to assess the feasibility of calling accurate *KIR* and *NKG2* genotypes when our dynamic diagnostic subregion approach is used with a comprehensive genomic allele reference library. From this data, we successfully identified all seventy-nine *KIR* and forty-eight *NKG2* alleles identified in our whole genome assemblies (FIGURE 3, 4). Given that the *KIR* and *NKG2* regions represent relatively small regions of the genome, we could infer the majority of genotypes despite having incomplete coverage of all alleles encoded within a given haplotype. For instance, if an animal encodes both a centromeric and telomeric gene from the same haplotype, we can assume the animal does not contain a recombinant haplotype with relatively high confidence. This highlights the power of using a population that has been fully defined at the genomic level. Still, the number of expected alleles identified within a given sample varied significantly. Most likely, this results from unequal loading of target capture products for sequencing between individual samples. The samples ranged from 22,458 to 93,963 *KIR* reads and 3,442 to 10,797 *NKG2* reads mapping to reference sequences, respectively. The exome capture reagent contained probes designed to capture rhesus macaque *KIR* sequences. Target capture efficiency for MCM samples could benefit from MCM-specific spike-in probes based on the genomic data presented here. Despite being a proof of concept, the data presented in Figures 3 and 4 demonstrates that multi-locus genotyping of complex immune gene families is at least feasible in a genetically-defined population like MCM.

**Figure 3.**
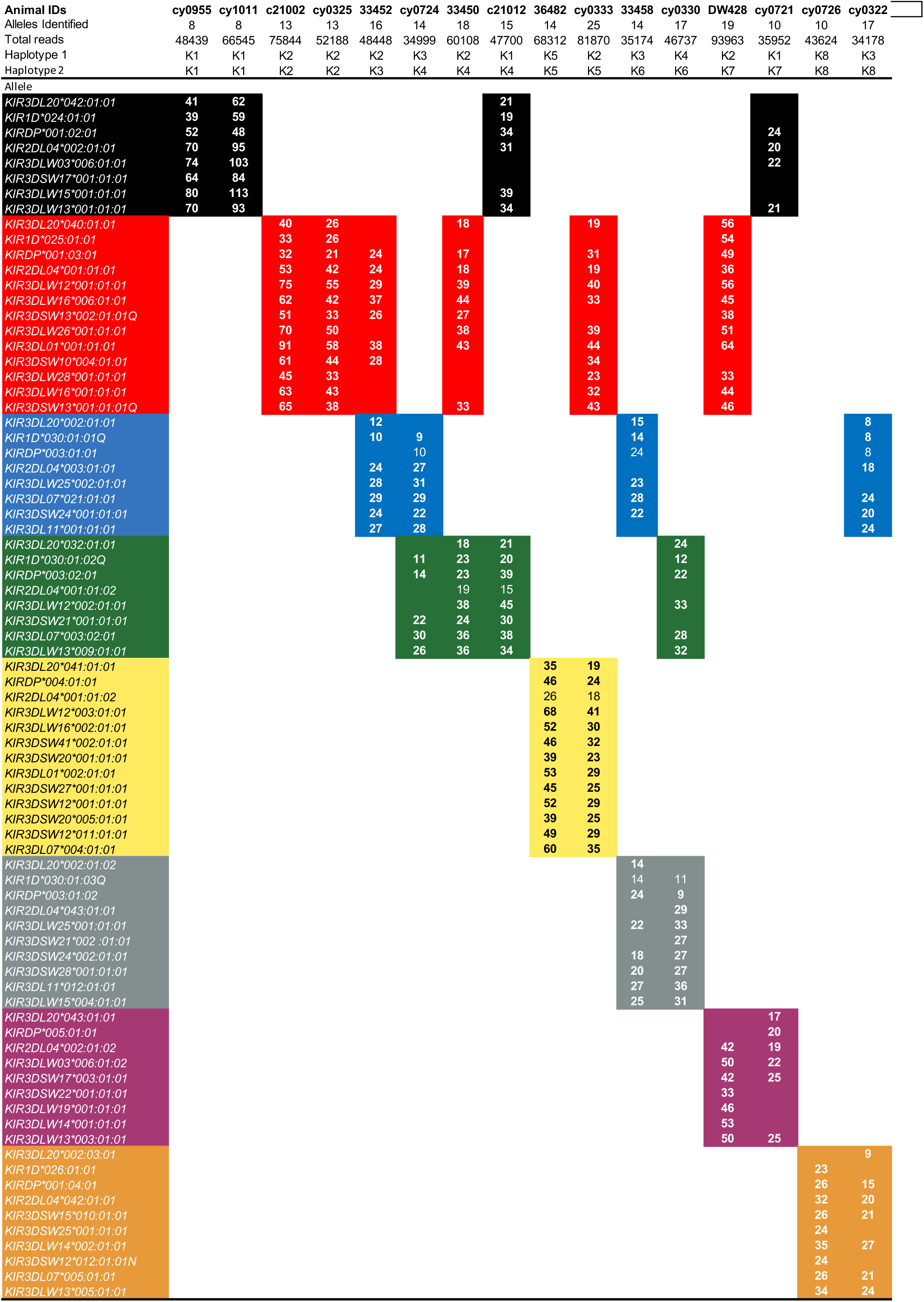
Exome genotypes for representative MCM. Alleles are displayed on the Y axis and are colored to denote the corresponding haplotype. The number of sequence reads mapped to algorithmically determined diagnostic sequences is presented within a single cell (see methods).

**Figure 4.**
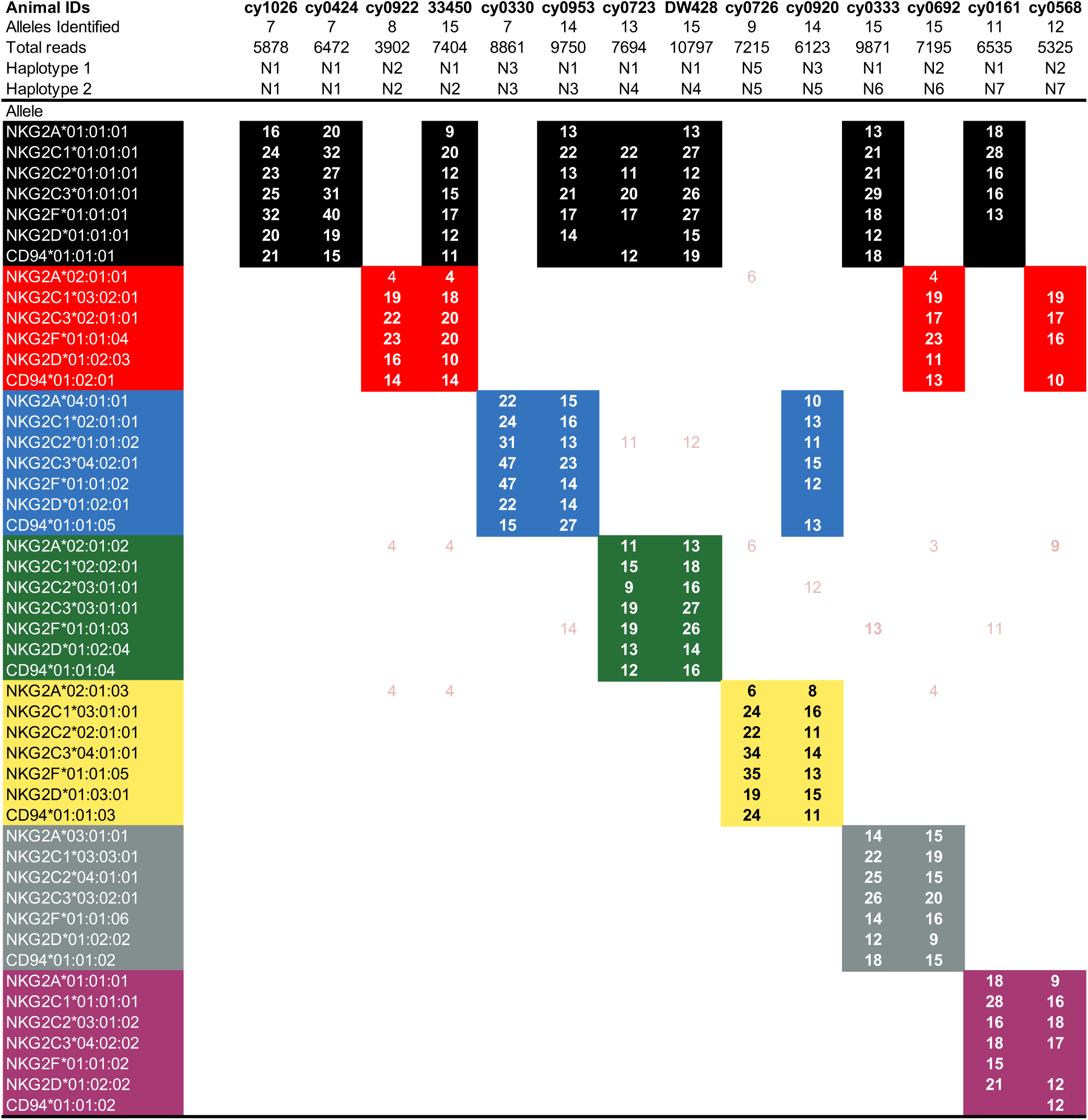
Exome genotypes for representative MCM. Alleles are displayed on the Y axis and are colored to denote the corresponding haplotype. The number of sequence reads mapped to diagnostic regions for each allele is presented within a single cell. Read counts displayed in pink represent presumptively miss-mapped reads due to near-identical diagnostic sequences between alleles. For instance, *NKG2C2:01:01:01* and *NKG2C2:01:01:02* are identical in sequence across exon intervals, and thus, samples will likely have reads that align to both alleles despite only having an N1 or N3.

*KIR* haplotype frequencies in the MCM population have been previously estimated using microsatellite markers (34). As this is the first characterization of the *NKG2* region within the species, no such frequency estimates exist. We calculated haplotype frequencies using data from fifty-nine unrelated animals out of our set of eighty-five exomes. As shown in Table II, the most common haplotype, N1, has a prevalence of 42%, followed by N2 (20%), N5 (14%), N3 (9%), N4 (6%), N7 (5%), N6 (2%), and recombinants (5%). Interestingly, both recombinant haplotypes seem to be a fusion of the N7 and N3 haplotypes, found in three, unrelated animals. This suggests the possibility of an eighth haplotype, akin to the N7 haplotype which is comprised entirely of a combination of *NKG2* variants from other haplotypes. Given the compact size of the *NKG2* region, recombination events are expected to be infrequent. However, our conclusions are based on a limited sample set, making these findings preliminary. A more comprehensive study with additional unrelated MCMs is necessary to accurately define the *NKG2* haplotype distribution within the MCM population.

## Discussion

Rapidly developing long-read sequencing technologies continue to improve the quality of whole-genome assemblies. This can be seen when comparing the continuity of the current cynomolgus macaque reference genome, MFA1912RKSv2, with its short-read predecessor (14). Despite these advancements, multigenic regions often remain inaccurately assembled and annotated. These inaccuracies are particularly visible within gene clusters integral to immune functions where relentless evolutionary pressure from persistent pathogen challenges has driven extensive diversification. Resolving these important immune-regulating gene clusters at the allelic level requires intricate manual curation. Recent efforts by multiple research teams have led to a comprehensive curation of immune regions in rhesus and cynomolgus macaques, shedding new light on the immune systems of these important biomedical model organisms (23, 33, 58).

Here, we have utilized long-read sequencing technology to conduct a comprehensive analysis of two key immune receptor families in the MCM population. Our study has identified fourteen novel and seventy nine total *KIR* alleles across eight ancestral haplotypes, thus capturing the entire spectrum of genetic variation present in this population. A similar examination of the *NKG2* region resulted in the assembly of seven unique haplotypes, the discovery of novel copy number variations, and the characterization of forty eight distinct alleles. Utilizing the high-resolution of these genomic assemblies, we have prototyped an approach for multi-locus genotyping using short-reads from whole exome datasets. Although this method requires further refinement, it has the potential to provide a cost-effective alternative to amplicon-based genotyping for multiple loci or expensive long-read genomic assembly techniques. Collectively, our findings offer an unparalleled insight into the genetic diversity of immune receptor families within an important non-human primate model, creating new opportunities to explore the relationship between genotype and phenotype for these critical immune components.

The evolving perspective amongst researchers suggests that natural killer (NK) cells play roles beyond controlling viruses and tumors; they also regulate inflammation and shape subsequent adaptive immune responses (17, 59). Extensive evidence has shown *KIR/MHC* allotype combinations impact the course of illnesses and infections such as preeclampsia (60), leukemia ((61–64)), human immunodeficiency virus (HIV) (64), and hepatitis C (65). Similarly, *NKG2/MHC-E* allotype combinations are implicated in differential responses to SARS-CoV-2 (66), HIV (62, 67, 68), and cytomegalovirus infections (63). These varied responses by NK cells are believed to stem from the influence of genetic sequences on the binding affinity between KIR and NKG2 receptors and their ligands (69). Despite progress in identifying KIR ligands in rhesus macaques (70, 71), most KIR in cynomolgus macaques remain without known ligands, partly due to a lack of specific immunohistochemical tools. However, the limited genetic variation within the MCM population may offer a more straightforward context for studying how genetics affects NK cell receptor/ligand interactions. The findings we present here provide a crucial basis for such foundational investigations of NK cell receptor biology.

Our analysis of KIR haplotypes revealed two groups defined by the number of *KIR3DS* genes in their telomeric segments. The first group (K1, K3, K4), parallels human group A *KIR* haplotypes in that they contain a single *KIR3DS* locus and minimal gene copy number variation (72). The second group of haplotypes (K6, K7, K8, K2, K5) have expanded copy numbers of *KIR3DL* and *KIR3DS* genes, resembling human group B haplotypes. The existence of both *KIR* group A and B haplotypes across human populations suggests they may be undergoing balancing selection (73). Given similar patterns in our data, macaques may be subject to a similar evolutionary balancing selection. This could potentially be tested using the whole genome data collected. For instance, population genetic assays such as the extended haplotype homozygosity analysis could measure slower decay in homozygosity for the *KIR* group A and B haplotypes, as compared to neutral expectations, providing evidence for ongoing balancing selection in these genomic regions (74). These types of statistical analyses of population dynamics will become increasingly useful as we resolve increasingly larger genomic space from the whole genome data reported here.

Recently, the non-classical class I molecule, MHC-AG, has been identified as a broadly recognized ligand of KIR3DS receptors in rhesus macaques (71). In humans, the engagement of HLA-C and soluble HLA-G with activating KIR2DS and KIR2DL4 on maternal NK cells located in the placental decidua triggers the secretion of pro-inflammatory and pro-angiogenic factors that support placental vascularization (60, 75). These pro-vascularization effects presumably decrease the risk of preeclampsia. Macaques lack orthologs for HLA-C and -G (76, 77). Instead, MHC-AG is theorized to have replaced their function due to its existence as both cell surface and soluble isoforms, as well as its broad expression in the placenta and amniotic membranes (16, 78–80). It’s plausible that interactions between KIR3DS and MHC-AG proteins may similarly influence placental development in macaques, especially given that at least one KIR3DS protein is encoded by every haplotype within the MCM population. This could be tested using large-scale genetic association studies that will be more approachable within MCM as cost-effective, multi-loci genotyping is refined. Furthermore, our comprehensive allele libraries open up the possibility for allele-level RNA expression analyses of placental and trophoblast tissues. Currently, immunohistochemical reagents are limited for macaques making KIR3DS and MHC-AG staining unavailable. Our genomic data can potentially aid in the creation of more KIR-specific antibodies, for instance, by immunization with KIR Fc fusion constructs.

The centromeric segments of *KIR* haplotypes in macaques are populated by lineage I *KIR* genes harboring less heterogeneity than their telomeric lineage II counterparts. We identified five novel *KIR3DL20* and two *KIR2DL04* CDS, elevating both genes to framework status within the population. Numerous studies have identified macaque *KIR2DL* transcripts that are analogous to human *KIR2DL5* and identical to rhesus *KIR3DL20* sequences. These *Mamu-KIR2DL5* transcripts have been considered to be alternatively spliced variants, or distinct alleles of *KIR3DL20*, where a single immunoglobulin domain is absent. The *KIR2DL5/KIR3DL20* locus has been proposed by some to potentially be an evolutionary stepping stone between the *KIR2DL* and *KIR3DL* genes (52). *KIR2DL04* in macaques is analogous to human *KIR2DL4* (54). It’s not possible to conclude that KIR2DL04 is necessary for fitness in macaques because previous sequencing studies have described KIR haplotypes without *KIR2DL04* in both rhesus and non-Mauritian cynomolgus macaques (33). In humans, KIR2DL4 is expressed by all natural killer cells, but unlike other KIR proteins, it is retained mostly within the endosomal compartment rather than being prominently expressed on the cell surface (81). Functionally, human KIR2DL4 can recognize HLA-G and initiate pro-inflammatory and anti-inflammatory responses, depending on the cellular context. Whether KIR2DL04 shares this functional significance in macaques is unknown, as its engagement with MHC-AG has not been documented. Still, low genetic diversity and its ubiquitous occurrence on most haplotypes could suggest positive selection within macaques. Further functional studies of lineage I genes will be necessary to elucidate their importance.

Our characterization of *NKG2* genomic haplotypes revealed the N2 haplotype has undergone a large deletion event, removing the entire *NKG2C2* gene from the chromosome. This is the first documentation of *NKG2C* copy number variation within macaques. We estimated the occurrence of this haplotype within the population at 20% based on exome sequencing datasets from unrelated animals. In humans, NKG2C+ NK cell subsets expand following acute human cytomegalovirus (HMCV) infection (82). This NKG2C+ population remains overrepresented in seroconverted patients and displays increased reactivity to HCMV-infected target cells in vitro and ex vivo after HCMV reactivation (83). Similar memory-like NK cell function has been observed in rhesus macaques following rhCMV and SIV infection (84). Recent research demonstrated that animals infected with rhesus cytomegalovirus (rhCMV) have expanded populations of NKG2C1+ and/or NKG2C2+ NK cells (85). Furthermore, single-cell RNAseq performed on rhCMV+ NK cells showed that *NKG2C2* is the most highly transcribed of the three *NKG2C* genes (85). Though we cannot say with certainty, our experience with polymorphic immune gene families in macaques has shown that observations in cynomolgus macaques often extend to rhesus macaques. If this *NKG2C2* deletion is also present within the rhesus population, this may have significant consequences in the context of rhCMV infection.

The pre-clinical vaccine strategy, utilizing strain 68-1 of rhCMV to express simian immunodeficiency virus (SIV) antigens, demonstrates the ability to induce effector memory T cell responses (86). These responses can restrict pathogenic SIV replication in approximately 50%–60% of vaccinated rhesus macaques (11, 87). The remaining ~45% of vaccinated yet unprotected monkeys demonstrate viral dynamics akin to those of unvaccinated controls. This binary phenotype is conserved in MCM when the vaccinations are performed with cyCMV vectors (11). Given the role of NKG2C in CMV immunity, *NKG2C2* copy number variation in macaques may provide at least a partial genetic basis for this intriguing phenotype. *NKG2* genotypes were not determined for animals used in these published vaccination experiments, as it has been assumed the *NKG2* region harbored very little genetic heterogeneity. Furthermore, the only NKG2C-specific antibody in rhesus macaques cannot distinguish between NKG2C1 and NKG2C2 proteins (11). The sequencing data presented here may prove useful in resolving this fascinating phenotype, whether it be through retrospective genotyping of animals used in these vaccination studies or by generating NKG2C1/2 specific antibodies using Fc fusion constructs similar to what’s performed when generating KIR allotype-specific reagents as previously mentioned (70).

In conclusion, our research provides an extensive genomic analysis of *KIR* and *NKG2* haplotypes within the MCM population. This thorough analysis has uncovered significant biological implications into crucial natural killer cell receptor genes. Leveraging high-quality reference assemblies, we have prototyped an economical genotyping approach, enabling extensive screening of these complex gene clusters at the colony level. We anticipate that the data and methodologies established in this study will form a crucial foundation for future biomedical research involving MCMs.

## Supporting information

Supplemental Table III

## Acknowledgements

We extended gratitude to Natasja de Groot and her expertise in providing the official IPD-NHKIR allele nomenclature for the KIR alleles reported in this study. We thank the anonymous reviewers for their insightful consideration of this manuscript and helpful suggestions to improve its content. We used the University of Wisconsin–Madison Biotechnology Center’s DNA Sequencing Facility (research resource identifier RRID:SCR_017759) to generate and sequence PacBio Sequel II HiFi libraries.

## Disclosures

The authors declare no competing interests.

## Data Availability

All raw and processed sequencing data generated in the study are publicly available through the National Center for Biotechnology Information (NCBI; https://www.ncbi.nlm.nih.gov/). The raw whole-genome ONT and PacBio HiFi data have been submitted to NCBI’s Sequence Read Archive (SRA; https://www.ncbi.nlm.nih.gov/sra) under BioProject PRJNA854973. Error-corrected genomic assemblies were submitted to GenBank under the accession numbers OR341018-OR341025 (KIR) and OR341105-OR341111 (NKG2). Extracted gDNA sequences for individual genes were submitted to GenBank. Associated accession genbank numbers can be viewed in supplemental tables II and III.

## Notes

This work was supported through contracts HHSN272201600007C and 75N93021C00006 from the National Institute of Allergy and Infectious Diseases of the National Institutes of Health. This work was also partly supported by the Office of Research Infrastructure Programs P51OD011106 awarded to the Wisconsin National Primate Research Center at the University of Wisconsin–Madison, and conducted in part at a facility constructed with support from Research Facilities Improvement Program grants RR15459-01 and RR020141-01.

https://www.ncbi.nlm.nih.gov/bioproject/?term=PRJNA854973

### Competing Interest Statement

The authors have declared no competing interest.

