## Supplemental Table I for "Complete genomic assembly of Mauritian cynomolgus macaque killer immunoglobulin-like receptor and natural killer group 2 haplotypes"

**Supplemental Table 1:** Genotypes and FASTQ statistics of thirteen sequenced animals

| sample | MHC | KIR | NKG2 | ONT reads KIR | ONT longest read KIR | PacBio reads KIR | PacBio longest read KIR | ONT reads NKG2 | ONT longest read NKG2 | PacBio reads NKG2 | PacBio longest read NKG2 |
|---|---|---|---|---|---|---|---|---|---|---|---|
| cy0161 | M2/M2 | K1/K1 | N1/N7 | 9520 | 420,616 bp | 1116 | 39,742 bp | 18689 | 508.639 bp | 2406 | 40.311 bp |
| cy0322 | M1/M1 | K3/K8 | N1/N4 | 3960 | 938,909 bp | 1420 | 40,498 bp | 7716 | 845,946 bp | 2989 | 43,031 bp |
| cy0325 | M1/M1 | K2/K2 | N1/N5 | 6538 | 413,461 bp | 1473 | 35,323 bp | 12529 | 423,355 bp | 3178 | 39,465 bp |
| cy0330 | M1/M3 | K4/K6 | N3/N3 | 4146 | 509,847 bp | 2009 | 42,052 bp | 8282 | 1,579,000 bp | 4201 | 45,862 bp |
| cy0333 | M3/M3 | K2/K5 | N1/N6 | 5548 | 431,005 bp | 2025 | 35,644 bp | 9568 | 491,527 bp | 3866 | 40,023 bp |
| cy0355 | M1/M5 | K4/K7 | N1/N2 | 2393 | 505,424 bp | 905 | 39,52 bp | 4999 | 664,884 bp | 2267 | 38,927 bp |
| cy0390 | M7/M7 | K3/K5 | N1/N3 | 1573 | 831,864 bp | 2054 | 39,706 bp | 3482 | 759,694 bp | 5014 | 42,86 bp |
| cy0424 | M4/M5 | K1/K3 REC | N1/N1 | 4419 | 532,42 bp | 2198 | 32,902 bp | 8728 | 861,247 bp | 4334 | 34,654 bp |
| cy0558 | M1/M3 | K7/K8 | N1/N1 | 2817 | 749,686 bp | 786 | 38,596 bp | 6228 | 778,774 bp | 2114 | 39,3 bp |
| cy0568 | M1/M3 | K2/K6 | N2/N7 | 3165 | 91,229 bp | 981 | 45,125 bp | 7066 | 158,677 bp | 2328 | 40,524 bp |
| cy0692 | M2/M4 | K3/K8 | N2/N6 | 3707 | 417,087 bp | 363 | 38,626 bp | 6787 | 412,564 bp | 1080 | 40,219 bp |
| cy0695 | M4/M4 | K2/K4 | N1/N7 | 5219 | 363,061 bp | 1296 | 41,196 bp | 9955 | 640,568 bp | 2771 | 44,042 bp |
| cy0973 | M6/M6 | K5/K7 | N1/N2 | 6972 | 674,889 bp | 1258 | 47,687 bp | 3648 | 745,565 bp | 2958 | 44,986 bp |
